# *Burkholderia* collagen-like protein 8, Bucl8, is a unique outer membrane component of a tetrapartite efflux pump in *Burkholderia pseudomallei* and *Burkholderia mallei*

**DOI:** 10.1101/2020.09.18.303271

**Authors:** Megan E Grund, Soo J Choi, Dudley H McNitt, Mariette Barbier, Gangqing Hu, Paul R LaSala, Christopher K. Cote, Rita Berisio, Slawomir Lukomski

**Author notes:** Corresponding author (SL).

## Abstract

Bacterial efflux pumps are an important pathogenicity trait because they extrude a variety of xenobiotics. Our laboratory previously identified *in silico Burkholderia* collagen-like protein 8 (Bucl8) in the Tier one select agents *Burkholderia pseudomallei* and *Burkholderia mallei*. We hypothesize that Bucl8, which contains two predicted tandem outer membrane efflux pump domains, is a component of a putative efflux pump. Unique to Bucl8, as compared to other outer membrane proteins, is the presence of an extended extracellular region containing a collagen-like (CL) domain and a non-collagenous C-terminus (Ct). Molecular modeling and circular dichroism spectroscopy with a recombinant protein, corresponding to this extracellular CL-Ct portion of Bucl8, demonstrated that it adopts a collagen triple helix, whereas functional assays screening for Bucl8 ligands identified binding to fibrinogen. Bioinformatic analysis of the *bucl8* gene locus revealed it resembles a classical efflux-pump operon. The *bucl8* gene is co-localized with downstream *fusCDE* genes encoding fusaric acid (FA) resistance, and with an upstream gene, designated as *fusR*, encoding a LysR-type transcriptional regulator. Using RT-qPCR, we defined the boundaries and transcriptional organization of the *fusR-bucl8-fusCDE* operon. We found exogenous FA induced *bucl8* transcription over 80-fold in *B. pseudomallei*, while deletion of the entire *bucl8* locus decreased the MIC of FA 4-fold in its isogenic mutant. We furthermore showed that the Bucl8 pump expressed in the heterologous *Escherichia coli* host confers FA resistance. On the contrary, the Bucl8 pump did not confer resistance to a panel of clinically-relevant antimicrobials in *Burkholderia* and *E. coli*. We finally demonstrated that deletion of the *bucl8*-locus drastically affects the growth of the mutant in L-broth. We determined that Bucl8 is a component of a novel tetrapartite efflux pump, which confers FA resistance, fibrinogen binding, and optimal growth.

**Author Summary:** *Burkholderia pseudomallei* and *Burkholderia mallei* are highly infectious and multidrug resistant bacteria that are classified by the National Institute of Allergy and Infectious Diseases as Tier one select agents partly due to the intrinsic multidrug resistance associated with expression of the efflux pumps. To date, only few efflux pumps predicted in *Burkholderia* spp. have been studied in detail. In the current study we introduce Bucl8, an outer membrane component of an unreported putative efflux pump with a unique extended extracellular portion that forms a collagen triple helix and binds fibrinogen. We demonstrate Bucl8’s role in fusaric acid resistance by defining its operon via bioinformatic and transcriptional analyses, as well as by employing loss-of-function and gain-of-function genetic approaches. Our studies also implicate the Bucl8-associated pump in metabolic and physiologic homeostasis. Understanding how Bucl8 efflux pump contributes to *Burkholderia* pathology will foster development of pump inhibitors targeting transport mechanism or identifying potential surface-exposed vaccine targets.

## Introduction

*Burkholderia pseudomallei* and *Burkholderia mallei* are Gram-negative bacteria that are the etiological agents of melioidosis and glanders, respectively [1]. Both pathogens are highly virulent and easily aerosolized, therefore they are classified as Tier one select agents by both the U.S. Department of Health and Human Services and the U.S. Department of Agriculture. In addition to being a biodefense concern, the bacteria are highly resistant to antibiotics and currently there is no licensed vaccine for either pathogen. Increasing global investigation into melioidosis has indicated that the disease may be more widespread than originally reported [2], and it has one of the highest disability-adjusted life years (DALY) of neglected tropical diseases at 4.6 million [3].

*B. pseudomallei* is a soil saprophyte that can infect humans, resulting in symptoms ranging from localized infections, including swelling or ulcerations, to systemic infections that can lead to septic shock [4].Treatment includes an extensive two-part chemotherapeutic regimen, most commonly using ceftazidime intravenously and then following it with an oral antibiotic eradication therapy of co-trimoxazole and doxycycline [5]. *B. mallei* is a clonal derivative of *B. pseudomallei* that has undergone significant genomic reduction and rearrangement. This genomic evolution is attributed to the species transition from being a soil saprophyte to an obligate host pathogen, selecting for genes advantageous for host-survival [6]. Glanders primarily affects equines, but can infect other livestock such as donkeys and goats. Although uncommon in humans, this zoonotic disease is often fatal if left untreated [4]. Symptoms typically affect the pulmonary system, including pneumonia and lung abscess, but may also present as cutaneous ulceration following direct inoculation.

Several classes of efflux pumps are expressed in multidrug resistant Gram-negative bacteria, such as *Pseudomonas aeruginosa*, *Acinetobacter baumannii*, and *Burkholderia spp*., and are at least partly responsible for their intrinsic antimicrobial resistance, including resistance-nodulation division (RND) efflux pumps [7]. *Burkholderia* are notorious for being resistant to an array of antibiotics, such as β lactams, aminoglycosides, tetracyclines, fluoroquinolones, macrolides, polymyxins, and trimethoprim [8], resulting in serious infections that are hard to treat. Bioinformatic analyses of the *B. pseudomallei* genomes have identified at least ten RND efflux pumps [9], although only three systems were characterized in more detail, *e.g.*, AmrAB-OprA, BpeAB-OprB, and BpeEF-OprC [10]; this gap in knowledge underscores a need for more studies of drug efflux pumps in *Burkholderia* [11]. Importantly, a large body of evidence indicates that efflux pumps also contribute to resistance to a variety of host-defense molecules, biofilm formation, regulation of quorum sensing and balanced metabolism, and overall pathogenesis [12], which further accentuate the importance of the efflux systems in bacteria.

Our previous studies have identified 13 novel *Burkholderia* collagen-like (CL) proteins (Bucl) containing collagen-like Gly-Xaa-Yaa or GXY repeats, as well as non-collagen domains, some of which had predicted functions [13]. Specifically, Bucl8 was predicted to be an outer membrane protein, containing tandem efflux pump OEP domains. Of the *Burkholderia* species tested, Bucl8 was present only in *B. pseudomallei* and *B. mallei*, although a homologous DNA sequence is present in *B. thailandensis.* Unique to Bucl8, as compared to typical outer membrane proteins, is an extended extracellular portion of unknown function that contains a presumed collagen-like domain, followed by a non-collagen C-terminal region. In addition, the collagen domain, which is broadly characterized as a stretch of repeating GXY motifs [14], in Bucl8 is composed of an uncommon repeating (Gly-Ala-Ser or GAS)_n_ collagen-like sequence.

Here, our objectives are to characterize the structure and function of the Bucl8-CL domain, define the *bucl8* locus, and identify substrates and potential function(s) of the putative Bucl8-associated efflux pump. We demonstrate that the collagen-like domain indeed adopts the characteristic collagen triple-helical structure. In addition, the recombinant extracellular portion of Bucl8 can bind to fibrinogen. We find that Bucl8 is the outer membrane component of an efflux pump responsible for fusaric acid (FA) resistance, a potent mycotoxin produced by *Fusarium* species that cohabitate the soil environment with *Burkholderia* [15, 16]. We further identify *bucl8*-associated genes encoding putative Bucl8-efflux-pump components. Transcripts of the *bucl8*-operon were upregulated in *B. pseudomallei* and *B. mallei* by exogenous FA, as well as by FA-derivative pHBA, which is involved in regulation of balanced metabolism in *E. coli*. FA resistance was diminished in a *B. pseudomallei* isogenic deletion mutant without the *bucl8* locus and could also be transferred to a FA-sensitive *E. coli* strain. Lastly, we found that the mutant grew at a significantly reduced rate, suggesting that under laboratory conditions the pump is important for the cell’s physiology. Here, we describe a previously unreported efflux pump with unique structure and functional implications in the biology of *B. pseudomallei* and *B. mallei species*.

## Materials and Methods

### Bacterial strains and growth

Two BSL2 *Burkholderia* strains exempt from the Select Agents list were used in this study: (i) *B. pseudomallei* strain Bp82 is an avirulent Δ*purM* mutant of strain 1026b [17], which was obtained from Christopher Cote (US AMRIID, Frederick, MD) and (ii) *B. mallei* CLH001 ΔtonBΔ *hcp1* mutant originates from the strain Bm ATCC23344 [18], which was obtained from Alfredo Torres (UTMB, Galveston, TX) (**Table 1**). Strain Bp82 was routinely grown in Luria broth-Miller (LBM) with shaking at 37°C and on Luria agar (LA) solid medium at 37°C. Strain CLH001 was grown under the same conditions, but the broth medium was supplemented with 4% glycerol. *E. coli* strains JM109 (Promega) and S17-1λpir/pLFX (*E. coli* Genetic Stock Center, Yale University) were cultured in LBM media and on LA. Antimicrobials were used in selective media and in susceptibility/ resistance assays, as described in the methods below.

**Table 1.** Bacterial strains and plasmids

| Strains and Plasmids |  | Description/ Characteristics | Source |
| --- | --- | --- | --- |
| *B. pseudomallei* | Bp 1026b (genomic DNA) | Blood culture from 29-year old female rice farmer with diabetes mellitus, Northeast Thailand, Sappasithiprasong hospital; 1993 | BEI Resources |
|  | Bp K96243 (genomic DNA) | Female diabetic patient- Khon Kaen hospital, Northeast Thailand; 1996 | BEI Resources |
|  | Bp82 | Attenuated 1026b strain with a partial deletion of the *purM* gene resulting in adenine and thiamine auxotrophy | USAMRIID, Frederick, MD |
| | Bp82$\Delta bucl8-fusE$ | Bucl8-pump deletion mutant | This study |
| *B. mallei* | CLH001 | Attenuated Bm ATCC23344 mutant with genes *tonB* (iron acquisition) and *hcp1* (type 6 secretory system structural protein) deleted | UTMB, Galveston, TX |
| *E. coli* | JM109 | Host strain; $\Delta endA1$, $\Delta recA1$, $\Delta lacZ$ gene | Promega |
|  | JM109::525 | JM109 with pSL525 plasmid containing the Bucl8-pump locus from Bp 1026b/Bp82 | This study |
|  | JM109::529 | JM109 with pSL529 plasmid containing the Bucl8-pump locus from Bp K96243 | This study |
|  | S17-1λpir/pLFX | Mobilization host | *E. coli* Genetic Stock Center, Yale University |
| Plasmids | pQE-30 | *E. coli* expression vector for proteins with N-terminal 6xHis-tag; T5 promoter; ampicillin resistance | Qiagen |
|  | pUC18T-mini-Tn7T-Tp | Mobilizable TpR mini-Tn7 vector; trimethoprim and ampicillin resistance | [19] |
|  | pMo130 | Mobilizable *E. coli* vector suicide in *Burkholderia* | [20] |
|  | pSL520 | pQE-30-based plasmid for expression of rBucI8-Ct protein | This study |
|  | pSL521 | pQE-30-based plasmid for expression of rBucI8-CL-Ct protein | This study |
|  | pSL522 | pMo130-based plasmid with *fusR* for generating chromosomal deletion of BucI8-pump. | This study |
|  | pSL524 | pMo130-based plasmid with *fusR* and *tar* for generating chromosomal deletion of BucI8-pump | This study |
|  | pSL525 | pUC18T-mini-Tn7T-Tp based plasmid with BucI8-pump locus of Bp 1026b/Bp82 | This study |
|  | pSL529 | pUC18T-mini-Tn7T-Tp based plasmid with BucI8-pump locus of Bp K96243 | This study |

### Bioinformatic analyses of the *bucl8* locus

#### Annotation of transcriptional and translational signals

The promoter regions of *fusR* and *bucl8* were defined by combining public transcriptome data and computational prediction. Briefly, strand-specific RNA-Seq data of *B. pseudomallei* [21] was downloaded from National Center for Biotechnology Information (NCBI) Sequence Read Archive (SRA) under BioProject accession PRJNA398168. The RNA-Seq read distribution across the genome was visualized by the UCSC genome browser [22], which includes a reference strain for 1106a. The genomic region spanning genes *fusR* to *tar* is highly similar between strain Bp 1106a and our target strain Bp 1026b (identity = 99.4%). The RNA-Seq reads were pooled and then mapped to the genome of strain 1106a using Bowtie2, which allows two base-pair mismatch [23]. The RNA-Seq read density at each genomic position was visualized by the UCSC genome browser [22] to determine putative transcription boundaries of *fusR* and *bucl8*. Sigma 70 promoters (−10 and −35) were predicted by BPROM [24]. Translation initiation sites (TISs) were predicted by TriTISA with default parameters [25]. The Shine-Dalgarno (SD) translation initiation signals were manually annotated within 20 bps upstream to TISs by considering “GGAG”, a SD consensus sequence annotated for *Burkholderia* [26]. The gene and protein designation were adopted according to Crutcher *et al*. 2017.

#### Prediction of FusR putative binding sites

The positions of the predicted FusR binding sites, a LysR-type transcriptional regulator, were determined using the University of Braunschweig Virtual Footprint Promoter analysis tool v3.0 [27]. Known LysR regulators were used as models to predict binding, including CysB, MetR, and OxyR from *E. coli*, GltC from *Bacillus subtilis*, and OxyR from *P. aeruginosa*. Standard settings were used to run the prediction (sensitivity = 0.8, core sensitivity = 0.9, and size = 5) on the 500-bp region upstream from the translational start site of *bucl8*.

### Genetic and molecular biology methods

#### Construction of an unmarked isogenic deletion mutant of bucl8 locus in Bp82

The chromosomal region in Bp82, encompassing genes *bucl8-fusCD-fusE*, was deleted using suicide plasmid pSL524 constructed in vector pMo130 (Addgene), as described previously [20].Two Bp82-DNA fragments of about 1 kb each were sequentially cloned within the multiple cloning site of pMo130: (i) pSL522 construct, containing *fusR* gene located upstream of *bucl8* was PCR-amplified with primers pSL522-ApaI-F and pSL522-HindIII-R, was cloned between *Apa*I-*Hind*III sites of the vector; and (ii) pSL524, containing *tar* gene located downstream of *fusE* was cloned at *Apa*I site, following amplification with primers pSL523-ApaI-2F and pSL523-ApaI-2R.

Plasmid pSL524 was introduced by conjugation into Bp82 via biparental mating with a donor strain *E. coli* S17-1λpir/pLFX::pSL524 on LA medium overnight. The mating bacteria were then scraped off and plated onto selective LA medium supplemented with 200 µg/mL kanamycin, to counter-select WT Bp82, and 50 µg/mL zeocin, to counter-select *E. coli*. Merodiploid colonies resulting from the single cross-over event, were sprayed with 0.45 M pyrocatechol (Sigma-Aldrich) to detect yellow transconjugants [20]. Several yellow colonies were streaked onto YT medium (10 mg/mL yeast extract, 10 mg/mL tryptone) containing 15% sucrose to force the excision of the *bucl8-fusE* locus and pMo130 sequence from Bp82 merodiploids. Colonies were grown for 48 hours. Successful excision produces deletion mutants as white colonies identified by spraying with pyrocatechol. White colonies were isolated and characterized by PCR and sequencing to confirm the deletion of the *bucl8-fusCD-fusE* genes.

#### Cloning of bucl8 locus in E. coli JM109

The cloning strategy was based on the genomic sequence of the Bp82 parent strain *B. pseudomallei* 1026b, which identified a ∼8.2-kb *Stu*I-*Stu*I fragment, encompassing the entire *fusR-bucl8-fusCD-fusE* locus. Bp82 gDNA was digested with *Stu*I and DNA species of about 8-10 kb were isolated from the gel and ligated to *Stu*I-cleaved vector pUC18T-mini-Tn7T-Tp (pUC18T-mini-Tn7T-Tp was a gift from Heath Damron, Addgene plasmid # 65024) [19].The *E. coli* JM109 transformants were isolated on a LA medium containing 100 μg/mL FA. Plasmid pSL525 was isolated from several colonies and analyzed by restriction digestion. Junctions between vector and insert sequences were sequenced to establish insert orientation. The presence of *fusR-bucl8-fusC-fusE* genes was verified by PCR and sequencing.

The plasmid construct pSL529, containing the *bucl8* sequence with extended CL region from Bp K96243, was also generated based on pSL525. An internal Bucl8 fragment from Bp K9264 (∼1.4-kb) was PCR-amplified (using primers Bucl8-1F and BurkhFusBCD-1R) and cloned between two unique sites in *bucl8*, *Xcm*I, and *Fse*I, of pSL525. *E. coli* JM109 transformants were isolated on a LA medium containing 100 μg/mL ampicillin. The plasmid sequence was verified as before.

#### Cloning, expression and purification of Bucl8-derived recombinant proteins

Two recombinant polypeptides, derived from the presumed extracellular portions of Bucl8 variant in Bp K96243, were generated for this study: (i) pSL521-encoded rBucl8-CL-Ct polypeptide, containing both the collagen-like region and the non-collagen C-terminal region and (ii) pSL520-encoded rBucl8-Ct, which only includes the C-terminal region.

For cloning, gBlocks (Integrated DNA Technologies) were designed, encoding two recombinant constructs (**Table S3**), as described [28]. gBlocks were used as templates to produce cloned DNA inserts using primers pSL521-F and pSL521-2R for pSL521 construct, and pSL520-F and pSL520-R for pSL520. gBlock DNA fragments were inserted between *Hind*III and *BamH*I sites of the pQE-30 vector, resulting in an N-terminal 6xHis-tag (Qiagen) for each construct and were then cloned into *E. coli* JM109. Plasmid constructs pSL520 and pSL521 were confirmed by sequencing (Primers pQE30-F, pQE30-2R).

For protein expression, *E. coli* JM109 with pSL520 or pSL521 constructs were grown in LBM plus 100 μg/mL ampicillin with shaking at 37C overnight, and then 10 mL cultures were used to inoculate 1 L batches of the same media. The protein expression was induced in cultures at OD_600_ ∼0.5 with 1 mM isopropyl β-d-1-thiogalactopyranoside for 3 hours and then bacterial cells were pelleted and frozen at −20°C overnight. Cell pellets were thawed and suspended in 10 mL of lysis buffer (50mM Tris buffer, 50mM NaCl, 2mM MgCl_2_, 2% Triton X-100, 10mM β-mercaptoethanol, 0.2 mg/mL lysozyme, 1 mL of EDTA free Protease inhibitor Mini tablets (Pierce), 1 mM PMSF, 10 µg/mL). The samples were vortexed, placed on ice for 20 minutes, and then centrifuged. The supernatants were applied onto affinity columns with HisPur^TM^ Cobalt Resins (Thermo Scientific) and purification was carried out according to manufacturer’s protocol. The eluted proteins were analyzed by 4-20% SDS-PAGE to assess the overall integrity and purity. The proteins were dialyzed in 25 mM HEPES and stored at −20°C.

### Ligand binding assay to rBucl8-CL-Ct and rBucl8-Ct

In the initial screening assay, binding of the rBucl8-CL-Ct to different extracellular matrix (ECM) ligands was assessed by ELISA [29]. Wells were coated overnight with 1 μg of each ligand dissolved in bicarbonate buffer: collagen type I and IV (Sigma), elastin (Sigma), fibrinogen (Enzyme Research), plasma fibronectin (Sigma), cellular fibronectin (Sigma), laminin (Gibco), and vitronectin (Sigma). Next, 1 μg per well of rBucl8-CL-Ct in TBS, 1% BSA was added and incubated for two hours at 37°C. Wells were washed with TBS and bound rBucl8-CL-Ct was detected with anti-6His-tag mouse mAb (Proteintech) in TBS-1% BSA and a secondary goat anti-mouse HRP-conjugated Ab (Jackson Immuno Research Laboratories Inc.); immunoreactivity was detected with ABTS substrate and measured spectrophotometrically at OD_415_. Data represent the mean ±SE of three independent experiments (n=3), each performed in triplicate wells. Concentration-dependent binding was assessed in a similar manner, however with varying concentrations (0-10 μM) of rBucl8-CL-Ct.

### Structural characterization of collagen-like domain

Homology modelling of the collagen-like (CL) region of Bucl8 was performed using the software MODELLER [30]. As a starting structure, we adopted the high-resolution structure of a collagen-like peptide (PDB code 1k6f) [31]. The Ct random coil region was generated using the Molefacture plugin of VMD [32]. Electrostatic potential surface was computed using the software Chimera [33].

Circular dichroism spectroscopy of rBucl8-derived polypeptides was performed as previously described [28]. Briefly, protein samples were dialyzed against 1x Dulbecco’s phosphate buffered saline, pH 7.4. CD spectra were taken with a Jasco 810 spectropolarimeter, in a thermostatically controlled cuvette, with a path length of 0.5 cm. Data were acquired at 10 nm per minute. Wavelength scans were performed from 240 nm to 190 nm at either 25°C or 50°C for unfolded triple helix in rBucl8-CL-Ct construct.

### Gene transcription by RT-qPCR

Duplicated bacterial cultures of Bp82 and CLH001 were grown in broth media at 37°C with shaking till early logarithmic phase (OD_600_ ∼0.4), then, FA was added to one of each culture at sub-inhibitory concentrations and incubated for one hour. Cultures were mixed with a 1:2 ratio of RNA Protect reagent, incubated for five minutes, then centrifuged and decanted. Pellets were suspended in lysing buffer (1.3 µg/µL proteinase K, 0.65 mg/mL lysozyme, TE; 10 mM Tris, 1mM EDTA, pH 7) and incubated for ten minutes. Total RNA was isolated using RNeasy Protect Bacteria Mini kit (Qiagen). TurboDNase enzyme [34] was used to remove traces of genomic DNA. RNA was either used immediately for cDNA synthesis or stored at −80°C for no more than one week. cDNA was generated using iScript cDNA synthesis kit (Bio-Rad #1708896).

RT-qPCR was performed using SsoAdvanced Universal SYBR Green Supermix (Bio-Rad), with primers listed in **Table S1**. Transcript levels were normalized to 16S rRNA [35]. Transcription fold change was calculated as relative to non-FA conditions, using the 2^−ΔΔCT^ method. Technical and experimental replicates were done in triplicate.

### Determination of antimicrobial susceptibility/resistance

#### Antimicrobial susceptibility by broth dilution method

Minimum inhibitory concentration (MIC) testing was performed in liquid and on solid media. Initially, Bp82 and CLH001 were grown overnight at 37°C with shaking to inoculate fresh media with varying concentrations of FA (32 µM to 8000 µM), as described [36]. The optical density was recorded after overnight incubation and colony forming units (CFU) were calculated after plating serially diluted samples on LA media.

#### Antimicrobial sensitivity on agar

Strains were also tested for growth on LA media supplemented with differing concentrations of antimicrobials. Bacterial cultures were grown to an optical density of ∼0.4 and plated on agar, and incubated at 37°C for 48 hours. The following concentrations of antimicrobials were used: fusaric acid (FA), 100-800 µg/mL [37]; para-hydroxybenzoic acid (pHBA), 0.5-2.5 mg/mL [38]; and chloramphenicol (CHL), 2-32 µg/mL serially diluted [39].

#### Antimicrobial susceptibility in clinical laboratory

Strains were tested with antimicrobials in a clinical laboratory using Thermo Sensititre GNX3F dehydrated 96-well plates (TREK Diagnostic Systems). Bacterial cultures were grown on LA medium and cells were emulsified in sterile water to turbidity of 0.5 McFarland. The suspension was then diluted in cation adjusted Mueller-Hinton broth with TES buffer before inoculation of 100uL (approximately 5×10^5^ CFU) into each antimicrobial test well. Plates were incubated for 24 hours at 34-36°C in a non-CO_2_ incubator. Results were read and interpreted based on manufacturer’s protocol and CLSI MIC interpretive guidelines [40]. Antimicrobials tested included: amikacin, doxycycline, gentamicin, minocycline, tobramycin, tigecycline, ciprofloxacin, trimethoprim/sulfamethoxazole, levofloxacin, aztreonam, imipenem, cefepime, meropenem, colistin, polymyxin, ceftazidime, cefotaxime, ampicillin/sulbactam, doripenem, piperacillin/tazobactam, ticarcillin/clavulanate.

Antimicrobial susceptibility was also assessed by disk diffusion using the following antimicrobials: ampicillin (Am 10), ciprofloxin (CIP 5), doxycycline (D 30), gentamycin (GM), trimethroprim/sulfamethoxazole (SXT), tetracycline (TE 30), tobramycin (NN 10), levofloxacin (LVX 5).

### Statistical analysis

Statistics were performed using GraphPad Prism software for two tailed paired Student’s *t* test, one way and two way ANOVA, pending the experiment. For gene expression of the *bucl8* operon in Bp82, statistical analysis was applied to log-transformed fold changes to account for the phenomena of heteroscedasticity. Significance was denoted at levels of \**p* ≤ 0.05, \*\**p* ≤ 0.01, \*\*\**p* ≤ 0.001. Error bars represent standard error measurements (SEM) with analyses based on three independent experimental repeats (n = 3), each performed in triplicate technical replicates, unless otherwise noted.

## Results

### Structural characterization of the extended extracellular domain of Bucl8 indicates triple helix formation

Previously, we had described the domain organization of the protein Bucl8, reported the coding sequence, and homology-modelled the structure of its periplasmic/outer membrane component, based on the structure of the outer membrane protein OprM of *P. aeruginosa* [13]. Each mature Bucl8 monomer is comprised of two tandem outer membrane efflux protein domains (OEP1 and OEP2), and a rare repetitive region consisting of glycine, alanine, and serine (GAS)_n_ triplet repeats, here denoted as the CL domain, which is followed by a non-collagen carboxyl-terminal (Ct) region (**Fig 1A**). Bucl8 is a homotrimeric molecule, which supports triple-helical structure of the extracellular CL-(GAS)_n_ domain, although, the specific (GAS)_n_ sequence has not been studied for triple helix formation. The number of consecutive (GAS)_n_ repeats present fluctuates between Bucl8 variants from different *B. pseudomallei* isolates. Analysis of ∼100 *bucl8* alleles showed (GAS)_n_ numbers ranging from 6 to 38 repeats (mode: 20). Notably, 21 consecutive GAS repeats characterize the Bucl8 of *B. pseudomallei* model strain K96243, while the Bucl8 variants of the strains utilized in this study have fewer (GAS)_n_ numbers, *e.g.*, Bp 1026b/Bp82 has six and *B. mallei* strain Bm ATCC 23344/CLH001 has eight. Following the CL-(GAS)_n_ domain is a Ct region of 72 amino acids that are conserved among *B. pseudomallei* and *B. mallei* strains. (**File S1**)

**Fig 1.**
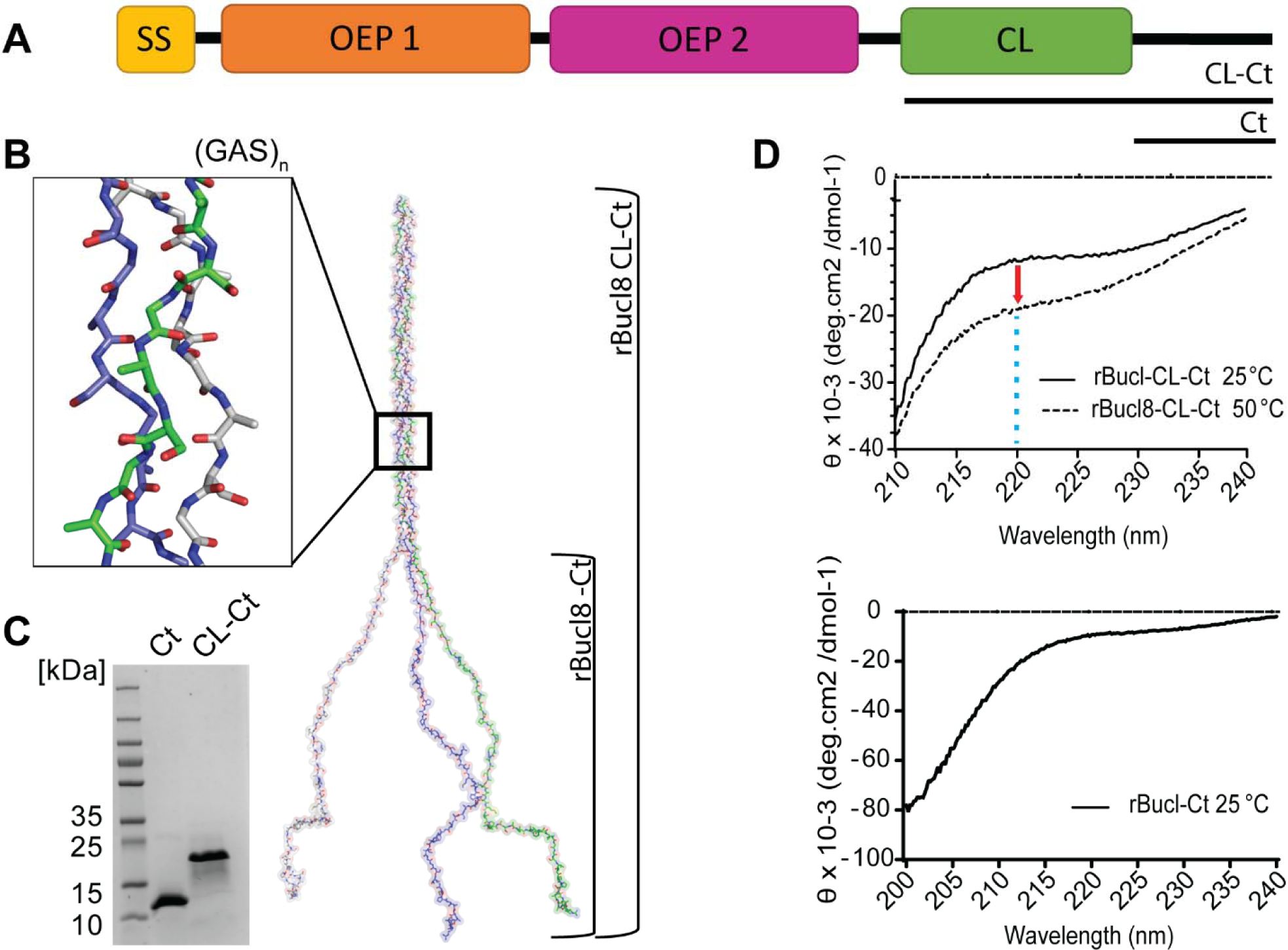
Structure analysis of extracellular component of the Bucl8 outer membrane protein. (A) Schematic organization of Bucl8 domain, including signal sequence (SS), outer membrane efflux protein domains 1 and 2 (OEP1, OEP2), collagen-like region (CL) and the C-terminus (Ct). (B) Structural modeling of Bucl8 extracellular region. Model depicts a homotrimeric polypeptide consisting of triple-helical CL domain of rBucl8-CL-Ct and unstructured C-terminus (rBucl8-Ct). The stick model in the inset depicts the triple helical fold of repeating (GAS)_n_ collagen sequence of Bucl8-CL. (C) 4-20% SDS-PAGE analysis of recombinant Bucl8-derived constructs. rBucl8-CL-Ct and rBucl8-Ct polypeptides were expressed in *E. coli* and purified via His-tag affinity chromatography. (D) Circular dichroism (CD) spectroscopy. (upper plot) Wavelength scans of rBucl8-CL-Ct were performed at 25°C (solid line) and 50°C (dashed line). A drop in molar ellipticity maximum at 220 nm (Θ_220_) is observed in the CD spectra, indicating the transition from triple-helical (25°C) to unfolded form (50°C). (bottom plot) CD spectrum of rBucl8-Ct at 25°C indicates an unstructured form.

Here, we homology-modelled a representative (GAS)_19_ sequence using the structure of the collagen peptide (PPG_10_) _3_ as a template (PDB code 1k6f, seqid 36%) [31] and the software MODELLER (**Fig 1B**). This structure formed a triple helix of about 163 Å in length. On its C-terminal end, the Ct domain of each chain is predicted by JPRED to be unfolded and was modeled in a random coil conformation (**Fig 1B**). Consistent with the sequence composition of the (GAS)_n_ repetitive domain, its electrostatic potential surface appears neutral, with only a few positive charges due to the presence of arginine residues in the unstructured Ct regions of the molecule (**Fig 1B**).

To experimentally validate this homology-modelled structure, two recombinant proteins, derived from the extracellular portion of Bucl8 variant in strain Bp K96243, were designed and expressed in *E. coli.* The construct rBucl8-CL-Ct includes the CL- (GAS)_19_ domain and adjacent unstructured C-terminus (Ct), while construct rBucl8-Ct encompasses the Ct region only. Both Bucl8-derived polypeptides migrate aberrantly in SDS-PAGE in relation to molecular weight standards, *e.g.*, rBucl8-CL-Ct of expected 11.7 kDa and rBucl8-Ct of 7.8 kDa (**Fig 1C**). Structural analysis of rBucl8-CL-Ct rendered at 25°C, using circular dichroism spectroscopy, confirmed a triple helical structure, demonstrated by a shallow peak at 220 nm (**Fig 1D**). As a control, denatured rBucl8-CL-Ct (50°C line) displayed a further-depressed peak at 220 nm that no longer held a triple-helical collagen structure. The 220 nm peak in rBucl8-CL-Ct is less pronounced when compared to typical triple helices formed by perfect GPP collagen repeats. This feature suggests the coexistence of both triple helix and random coil structures and/or the contribution of the non-collagen Ct region to the spectrum; such effects on CD spectra were previously reported for streptococcal collagen-like rScl constructs [41]. Additionally, the rBucl8-Ct structure was also analyzed by circular dichroism spectroscopy. The absence of ellipticity maxima and/or minima of known structures, *e.g.*, α-helices or β-strands [42], indicates an unstructured protein (**Fig 1D**). Altogether, using *in silico* modeling and experimental CD spectroscopic analyses of the representative recombinant protein, we demonstrated that repeating (GAS)_n_ of the predicted Bucl8-CL region from *B. pseudomallei* and *B. mallei* can form a stable collagen triple helix; to our knowledge, this is the first such demonstration obtained for the unusual repeating (GAS)_n_ collagen-like sequence.

Bacterial proteins harboring CL domains from diverse genera have been demonstrated to bind ligands, including extracellular matrix proteins (ECM), and have been shown to participate in pathogenesis [43–45]. Here, we screened several human compounds by ELISA to ascertain a potential ligand binding function of Bucl8’s extracellular region, rBucl8-CL-Ct; ligands included fibrinogen, collagen-I and IV, elastin, plasma and cellular fibronectin, and vitronectin. Of the ligands tested, rBucl8-CL-Ct construct showed significant binding to fibrinogen, but not to collagen I and elastin (**Fig 2A**), while binding to other ligands tested was also not significant (not shown). rBucl8-CL-Ct binding to fibrinogen-coated wells was concentration-dependent in contrast to control BSA-coated wells. In addition, rBucl8-Ct construct showed limited level of binding to fibrinogen in this assay (**Fig 2B**).

**Fig 2.**
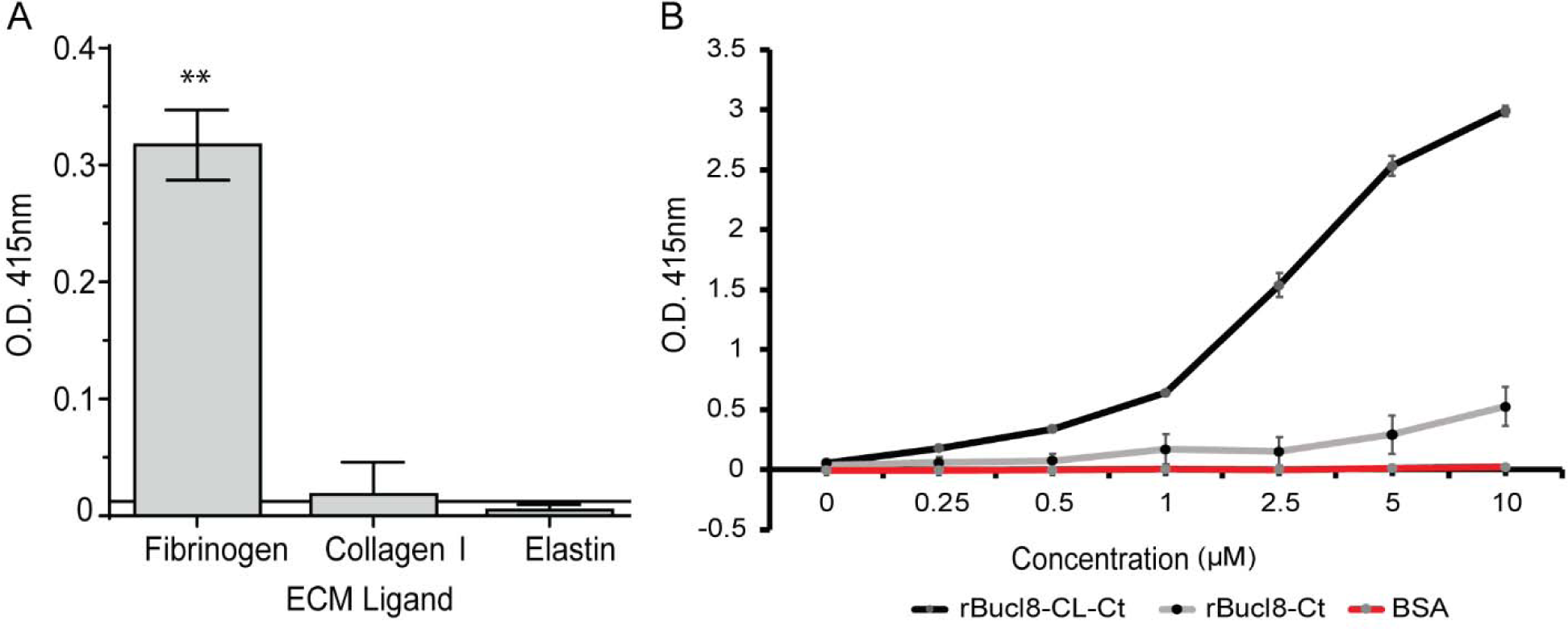
Binding of rBucl8-derived constructs to extracellular matrix proteins. Screening assay for rBucl8-CL-Ct binding to extracellular matrix proteins. Ligand binding was tested by ELISA; representative examples of rBucl8-CL-Ct-binding-positive and binding-negative ligands are shown. rBucl8-CL-Ct binding was compared statistically with binding to BSA-coated wells plus two standard deviations; Student’s *t*-test, **p ≤ 0.01. (B) Concentration-dependent binding of rBucl8-CL-Ct and rBucl8-Ct to fibrinogen. Wells were coated with fibrinogen and either recombinant Bucl8-derived protein was added at increasing concentrations. Data represents the mean ±SEM of three independent experiments (n=3), each performed in triplicate wells. Binding was detected with an anti-His-tag mAb.

### Identification of *bucl8* operon in *Burkholderia pseudomallei* and *Burkholderia mallei*

Previously, we identified two tandem outer-membrane-efflux-protein (OEP; PF02321) domains in Bucl8 [13], leading to the current hypothesis that Bucl8 is the outer membrane component of an efflux pump. Genes encoding efflux pumps are often clustered in operons that are controlled in *cis* by transcriptional regulators, such as MexR of *P. aeruginosa* and AmrR of *B. pseudomallei* [46–48]. For this reason, we examined the genes surrounding *bucl8*, which are described in **Table 2** and depicted in **Fig 3A**. The locus contains additional efflux-pump associated genes, annotated in the NCBI database to be involved in fusaric acid (FA) resistance, which we designated here as ‘*fus*’, as previously proposed [37]. In agreement with genomic annotations, we recognize that Bucl8 is an outer membrane lipoprotein with a lipid moiety attached via the N-terminal Cys residue of the mature protein (**Fig 3B**; residue No. 24). In the genome of *B. pseudomallei* 1026b, downstream of *bucl8* (OMP; 594 aa) are: *fusC*, presumably encoding the inner membrane protein of the pump (IMP; 733 aa), *fusD*, encoding a small protein with domain of unknown function (DUF; 67 aa), and *fusE* encoding the periplasmic adaptor protein (PAP; 293 aa). The ATG start codon of *fusD* overlaps with a stop TGA codon of *fusC*. The direction of the next downstream gene, *tar*, is opposite to *bucl8*-*fusCDE* and was presumed by definition to be outside of this operon. Flanking the locus at the 5’ end of *bucl8* is a divergently-oriented gene, encoding a LysR-type transcriptional regulator (LysR; 313 aa) [49], designated here as *fusR*. The proximity and opposite orientation of *fusR* gene in relation to the *bucl8-fusE* genes resembled the typical gene organization described in tripartite efflux pumps with LysR-type regulators; therefore, we hypothesized *bucl8* transcription to be regulated by the *fusR* product. Using predictive software and analysis of transcriptome data, the promoters, transcription initiation sites (TIS), and FusR binding sites were identified in the intergenic region between *fusR* and *bucl8* (**Fig 3B**). FusR was predicted to have four binding sites, depicted in the green boxes that overlap with the *bucl8* −10 and −35 sites. The consensus sequence for *B. pseudomallei* is “GGAG”, according to the ProTISA database [25], which matches *bucl8*’s predicted Shine-Dalgarno sequence. Thus, *fusR-bucl8-fusCD-fusE* constitute a regulon, likely involved in FA resistance. The *bucl8* locus was also conserved in Bp strain K96243 and Bm ATTC 23344; however, transcriptional units of *bucl8-fusE* were on the positive strand in the genome of K96243 strain, and on the negative strand in Bp 1026b and Bm ATTC 23344 (**Table 2**).

**Fig 3.**
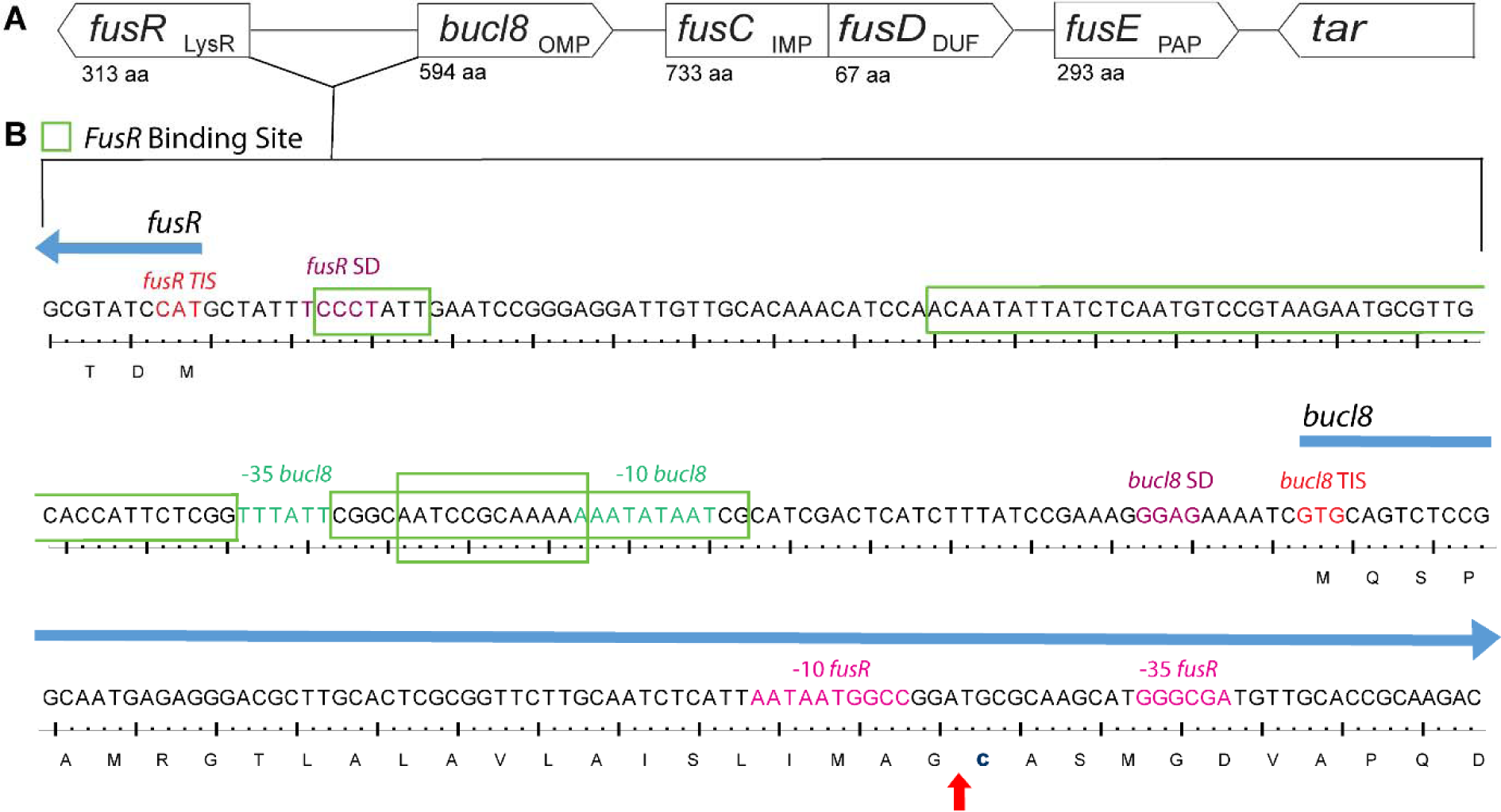
Chromosomal locus surrounding *bucl8* gene in *B. pseudomallei* and *B. mallei*. (A) Schematic of *bucl8*-associated locus with presumed protein function (subscript) and amino acid length (aa). Upstream of *bucl8* is gene *fusR*, while downstream are genes *fusCD* and *fusE.* Flanking the *bucl8* operon is unrelated downstream gene *tar*. LysR, LysR-type transcriptional regulator; OMP, Outer membrane protein; IMP, Inner membrane protein; DUF, Domain of unknown function; and PAP, periplasmic adaptor protein. (B) Regulatory intergenic region between *fusR* and *bucl8*. Both nucleotide and translated sequence are shown. Red arrow indicates cleavage site between the signal peptide and N-terminal cysteine linker (bolded).

**Table 2.** Genes and associated identification numbers of *bucl8* locus.

| Gene | Product Annotation | Bp 1026b |  |  | Bp K96243 |  |  | Bm ATTC 23344 |  |  |
| --- | --- | --- | --- | --- | --- | --- | --- | --- | --- | --- |
|  |  | Locus tag | Protein ID | Genomic position | Locus tag | Protein ID | Genomic position | Locus tag | Protein ID | Genomic position |
| *fusR* | Transcriptional regulator | BP1026B_I1940 | AFI66557.1 | 2150545-2151486 | BPS_R S10485 | WP_0045 34689.1 | 2345922-2346863 | BMA_RS 04430 | WP_0041 91155.1 | 987878-988819 |
| *bucI8* | RND efflux system, outer membrane lipoprotein, NodT family protein | BP1026B_I1941 | AFI66559.1 | 2151644-2153428 | BPS_R S10490 | WP_1624 86666.1 | 2347036-2348973 | BMA_RS 04425 | WP_0249 00385.1 | 985939-987705 |
| *fusC* | Fusaric acid resistance protein | BP1026B_I1942 | AFI66560.1 | 2153445-2155646 | BPS_R S10495 | WP_0099 37757.1 | 2348990-2351191 | BMA_RS 04420 | WP_0041 92976.1 | 983721-985922 |
| *fusD* | Hypothetical protein | BP1026B_I1943 | AFI66561.1 | 2155643-2155846 | BPS_R S10500 | WP_0041 91885.1 | 2351188-2351391 | BMA_RS 04415 | WP_0041 91885.1 | 983521-983724 |
| *fusE* | Fusaric acid resistance protein *fusE* | BP1026B_I1944 | AFI66562.1 | 2155860-2156741 | BPS_R S10505 | WP_0045 34908.1 | 2351405-2352286 | BMA_RS 04410 | WP_0041 91342.1 | 982626-983507 |
| *tar* | Methyl-accepting chemotaxis protein | BP1026B_I1945 | AFI66563.1 | 2157092-2158768 | BPS_R S10510 | WP_0041 96082.1 | 2352657-2354333 | BMA_RS 04405 | WP_0041 96082.1 | 980610-982286 |

Data were retrieved from NCBI for *B. pseudomallei* 1026b, *B. pseudomallei* K96243, and *B. mallei* ATTC 23344 reference genomes.

### Fusaric acid increases relative expression of *bucl8*-operon transcripts

We identified a conserved operon associated with the *bucl8* gene that was present in all *B. pseudomallei* and *B. mallei* genomes analyzed, including the mutant strains Bp82 and CLH001 used in this study, and had similarity to genes encoding FA resistance found in other Gram-negative bacteria [15, 16, 37]. We consequently tested the predicted FA substrate as a transcriptional inducer for genes associated with the Bucl8-efflux pump. We first examined MICs for FA resistance in both *B. pseudomallei* and *B. mallei* strains using a broth dilution method in the range of 32 µM FA to 8000 µM, which was based on an earlier induction data employing GFP reporter construct in *P. putida* [36]. Here, we established the FA-MIC for Bp82 as 4000 µM (716 µg/mL) and 250 µM (44 µg/mL) for CLH001.

Sub-inhibitory concentrations of FA, *e.g.*, 1000 µM for Bp82 and 60 µM for CLH001 that did not inhibit the growth rates were used in subsequent induction experiments (**Fig 4A**). Total RNA was isolated from the cultures of Bp82 and CLH001 that were either non-treated or treated with FA (1000 µM or 60 µM, accordingly) at OD_600_ ∼0.4 for one hour. Both *fusR* and *bucl8* genes were expressed in non-treated cultures at basal levels, but transcription of *bucl8* in Bp82 was significantly induced with FA by an average 82-fold change in relative expression and a 20-fold change of *fusR*, using 2^ΔΔ^Ct calculations (**Fig 4B**). CLH001 also demonstrated about a four-fold increase for *fusR* and *bucl8* when induced with 60 µM FA (**Fig 4C**), although this change is comparatively lower than that recorded in FA-induced Bp82.

**Fig 4.**
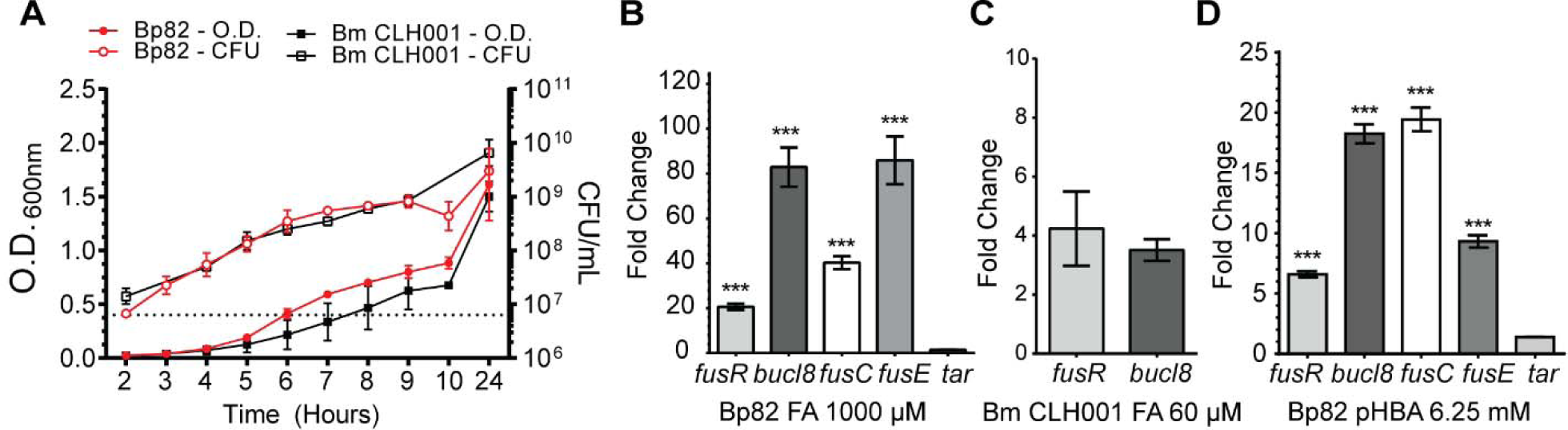
Effect of FA and pHBA on gene transcription within *fusR-bucl8-fusCD-fusE* operon. (A) Growth curves of *B. pseudomallei* strain Bp82 and *B. mallei* strain CLH001. Cultures were grown in strain-specific broth and optical density (O.D.) and colony forming units (CFU) were recorded. Dotted line represents OD of 0.4. Error bars represent ±SEM. (B-D) RT-qPCR was performed on RNA samples isolated from cultures of the indicated strain, untreated and treated with substrate, at an OD_600_ of ∼0.4 for 1 hour. Graph shows fold change of relative gene expression compared to untreated cultures and normalized to transcription of 16S rRNA gene. Technical and experimental replicates were done in triplicate. One-way ANOVA with Tukey’s multiple comparison test of the log_10_ –transformed fold change. Significance shown is in comparison to *tar*; ***p < 0.001. Error bars represent ±SEM. (B) Transcription activation of *fusR-bucl8-fusCD-fusE* genes in Bp82 with 1000 µM FA. The downstream *tar* gene is assumed outside of the *fusR-bucl8* operon. (C) Transcription activation of *fusR* and *bucl8* in CLH001 with 60 µM FA. (D) Transcription activation of Bucl8 regulon in Bp82 with 6.25 mM pHBA.

In a following experiment we confirmed the boundaries of the *fusR*-*bucl8* operon by RT-qPCR. Results show that transcription levels of *fusR*-*bucl8-fusC-fusE* were all significantly upregulated in samples treated with FA, compared to non-treated controls (*fusR* = 20-fold ± 1.37; *bucl8* = 82-fold ± 8.73; *fusC* = 40-fold ± 2.84; *fusE* = 86-fold ± 10.65; **Fig 4B**). In contrast, the expression change of *tar* was significantly lower than genes from the *fusR*-*bucl8-fusC-fusE* operon and the associated regulatory gene *fusR* (1.5-fold ± 0.03. One-way ANOVA with Tukey’s multiple comparison test of the log_10_-transformed fold change; ***p < 0.001 for all genes compared to *tar*). This is the first demonstration of FA-inducible efflux pump in *B. pseudomallei* and *B. mallei*.

### A structural analog of fusaric acid pHBA induces pump expression

Previous work reported that FusC-containing FA-exuding pumps were phylogenetically related to the aromatic carboxylic acid (AaeB) pumps, although it was unknown whether AaeB systems extrude FA [37]. Notably, studies in *E. coli* show that regulated concentrations of an FA-derivative, para-hydroxybenzoic acid (pHBA), inside bacterial cells is important for balanced metabolism of the aromatic carboxylic acids [38]. Thus, we hypothesized pHBA would also increase the relative expression of the *bucl8* operon as FA did. Broth cultures of Bp82Δ*bucl8-fusE* were induced with the sub-inhibitory concentration of 6.25 mM (863 µg/mL) pHBA and compared to non-treated cultures. RT-qPCR data showed in Bp82 pHBA induced a 7-fold ± 0.26 change in *fusR*, an 18-fold ± 0.78 change in *bucl8*, a 19-fold ± 0.98 change in *fusC*, and a 9-fold ± 0.52 change in *fusE.* Transcription of *tar* was not significantly affected (1.4 fold ± 0.006 change; One-way ANOVA with Tukey’s multiple comparison test of the log_10_-transformed fold change; ***p < 0.001 for all genes compared to *tar*) (**Fig 4D**). Evidence that aromatic carboxylic acids can induce transcription of this pump may help elucidate the broader function of Bucl8-associated pump in *B. pseudomallei and B. mallei*.

### Deletion of and complementation with the Bucl8-pump affect sensitivity and resistance to FA and pHBA

In order to demonstrate the function of the Bucl8-pump in various physiological roles, we used a genetic approach by generating two strains for assessing (i) loss-of-function and (ii) gain-of function. For loss-of-function, we made an isogenic Bp82 mutant harboring chromosomal deletion of *bucl8-fusCD-fusE* segment, as described [20]. Plasmid pSL524 (**Table 1**) was constructed in the *E. coli* vector pMo130, which is suicidal in *Burkholderia*, to generate an unmarked deletion mutant (**Fig 5A**). Construct pSL524, carrying upstream and downstream sequences flanking *bucl8* locus was transferred to *B. pseudomallei* Bp82 via biparental mating. Deletion was achieved in a two-step insertion/excision process, as detailed in Materials and Methods section. Successful deletion of the *bucl8-fusCD-fusE* segment from the chromosome was confirmed by PCR (**Fig 5B**) and sequencing. We did not delete the *fusR* gene on purpose to avoid a possible global regulatory effect associated with unknown FusR function.

**Fig 5.**
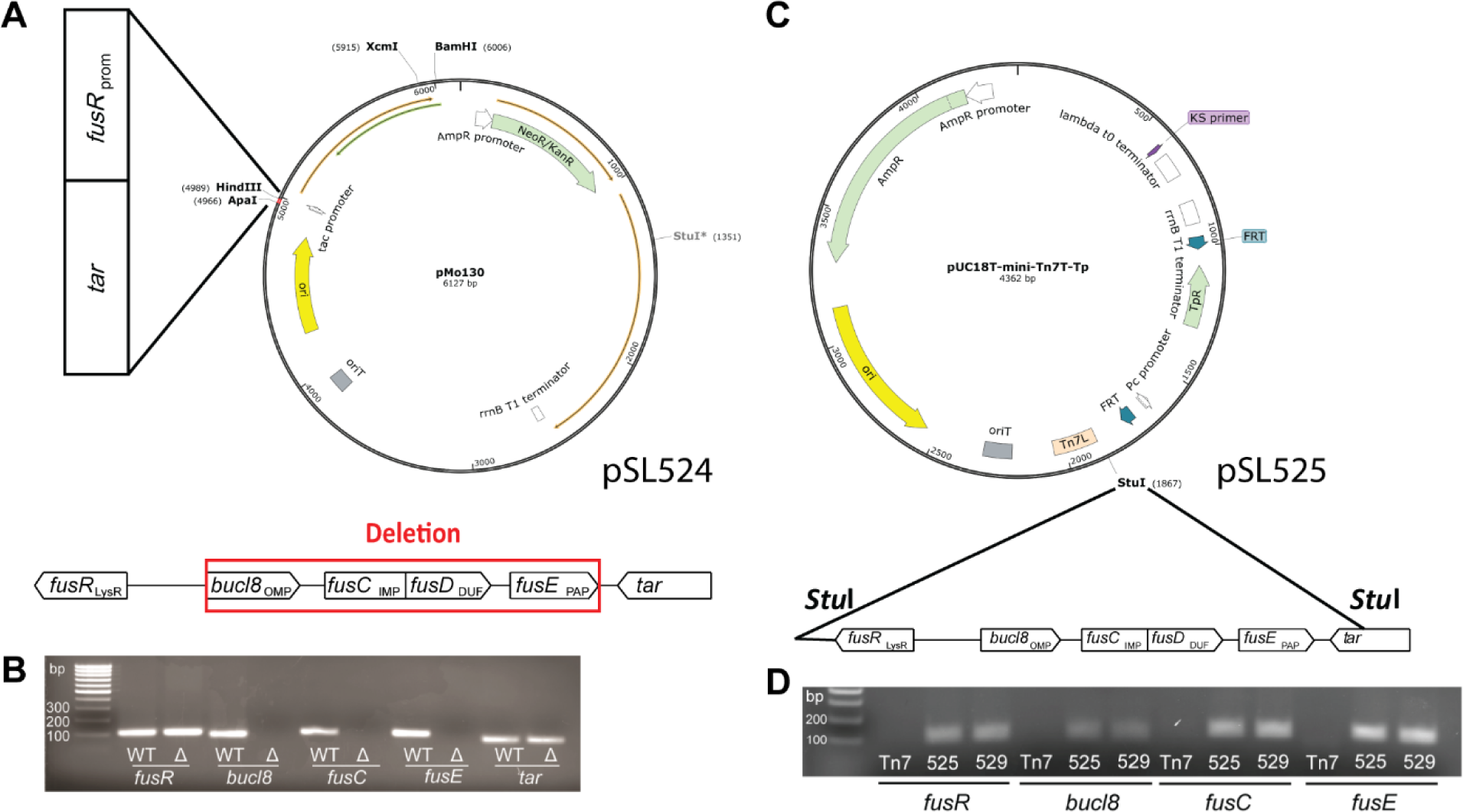
Construction of an unmarked Bucl8-pump deletion mutant and cloning into a heterogenetic host. (A) Strategy for generating an unmarked Bucl8-pump deletion mutant. Construction of the suicide plasmid construct pSL524. Vector pMo130, which is suicide in *Burkholderia*, was used to generate pSL524 plasmid construct for mutagenesis. *Hind*III and *Apa*I sites were utilized to clone flanking regions containing *fusR* and *tar* sequences to delete the *bucl8-fusE* coding region, depicted below. (B) Analysis of the *bucl8-fusE* deletion mutant of Bp82 by PCR. The presence of *bucl8-fusE* genes was tested in the genomic DNA isolated from wild type Bp82 (WT) and Bp82 *bucl8-fusE* mutant (Δ). (C) Cloning of the Bucl8-pump locus for *in-trans* complementation in *E. coli*. Vector pUCT18T-mini-Tn7T-Tp was used for cloning of an 8.2-kb genomic Bp82 fragment, flanked by *Stu*I sites, encompassing *bucl8* locus. (D) Characterization of the pSL525 and pSL529 constructs. The presence of *fusR-fusE* genes on pSL525 and pSL529 plasmids was tested by PCR. PCR products shown in B and D were analyzed on 1.3% agarose gel.

To exhibit gain-of function, we complemented a heterologous *E. coli* host *in-trans* with a plasmid construct pSL525 (**Table 1**) harboring the whole *bucl8* locus, generated in a mini-transposon vector pUC18T-mini-Tn7T-Tp, as depicted in **Fig 5C**. JM109::525 transformants were selected on agar containing 100 μg/mL FA and cloning was verified by PCR (**Fig 5D**) and sequencing. Since Bp82 represents the 1026b strain harboring Bucl8 variant with (GAS)_6_ repeats in the CL region, we made an additional construct, pSL529, that contains (GAS)_21_ repeats, to represent the majority of *B. pseudomallei* strains, by extending the number of GAS triplets in pSL525.

MICs were determined for bacterial growth on LA plates containing FA or pHBA chemicals, ranging from 0 to 800 μg/mL FA and 500-2,500 μg/mL pHBA (**Fig 6A**). There was a 4-fold decrease in MIC to FA from 400 µg/mL to 100 μg/mL recorded for Bp82Δ*bucl8-fusE* mutant compared to the parental Bp82 strain. A similar effect was observed for pHBA; the MIC for Bp82 was 1500 μg/mL which decreased to 1000 μg/mL in the mutant. A 12-fold increased MIC on the LA medium with FA was recorded in *E. coli* JM109::525 and JM109::529 (MIC = 300 µg/mL) compared with the JM109 (MIC = 25 µg/mL) recipient. Interestingly, complementation with Bucl8-pump, however, did not increase the MIC for pHBA above 1000 μg/mL for JM109::525 or JM109:529 strains.

**Fig 6.**
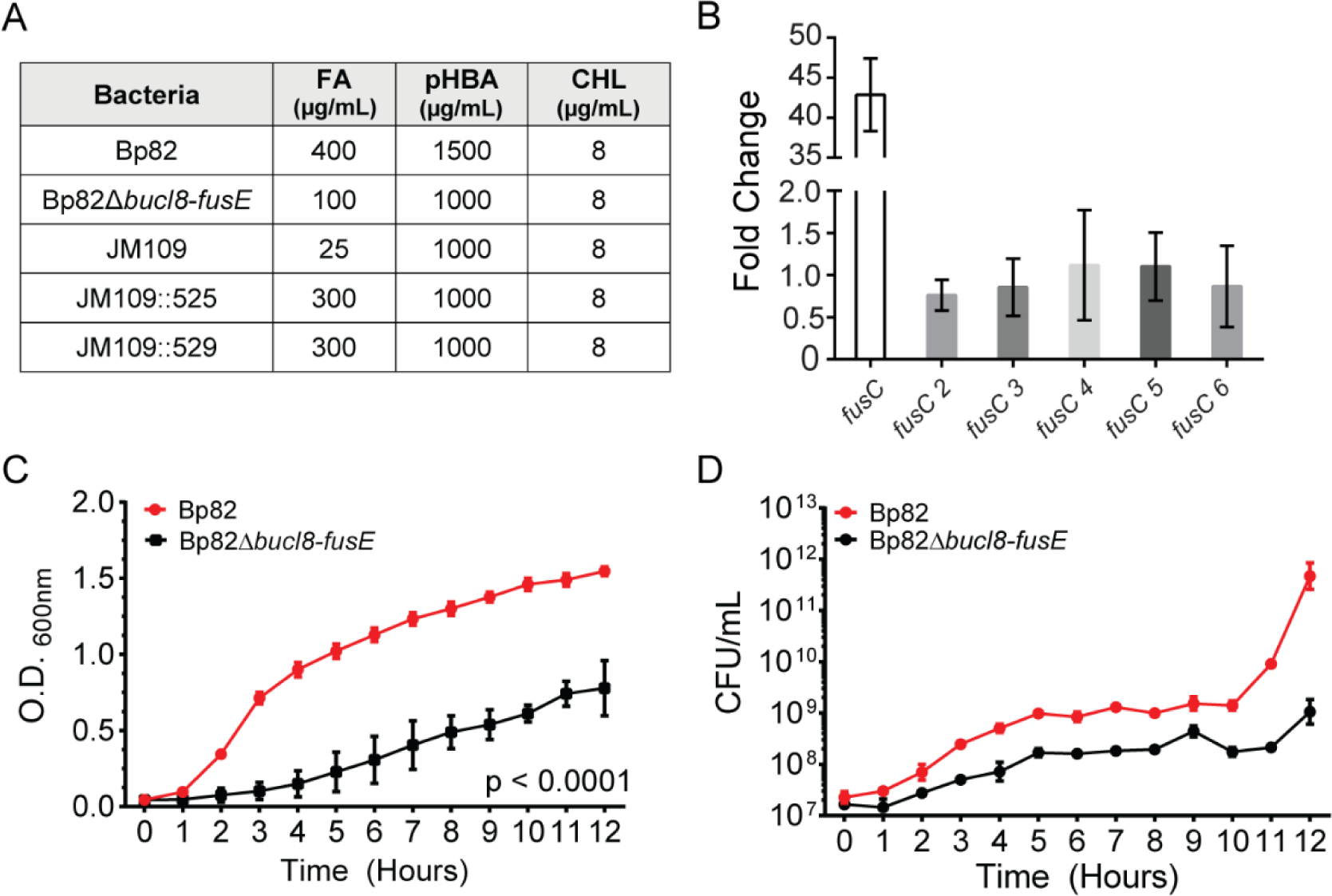
Analysis of loss-of-function and gain-of-function associated with chromosomal deletion and *in-trans* complementation of Bucl8-pump components. (A) Changes in sensitivity/resistance patterns in bacterial strains. MIC was determined by plating bacteria on LA containing each substrate. FA, fusaric acid; pHBA, para-hydroxybenzoic acid; CHL, chloramphenicol. (B) Relative expression of *fusC* genes. RT-qPCR was performed on total mRNA isolated from non-treated and FA-treated (1000 µM, 1 hour) Bp82 cultures (OD_600_ ∼0.4). Graph shows fold change of relative gene expression compared to untreated cultures and normalized to 16S rRNA. Technical and experimental replicates were done in triplicate. (C-D) Effect of chromosomal deletion on growth. Parental strain Bp82 and its *bucl8-fusE* deletion mutant (Bp82D*bucl8-fusE*) were grown in LBM broth at 37°C with shaking. Changes in OD_600_ (C) were recorded and CFU numbers (D) by plating on LA medium every hour. Data represents the average of three biological replicates. 2-way AVOVA with Tukey multiple comparison test, ***p < 0.001. Error bars represent ±SEM.

Although deletion of the Bucl8 pump resulted in a drastically decreased MIC, Bp82Δ*bucl8-fusE* mutant still maintained residual level of FA resistance (100 μg/mL).Therefore, we hypothesized that additional proteins annotated as FusC are contributing to the remaining FA resistance recorded in the Bp82Δ*bucl8-fusE* mutant. Within Bp 1026b and K96243 genomes, there are six genes present that are annotated as FusC-type proteins (Pfam #PF04632), including the protein arbitrarily designated as FusC, which is associated with Bucl8, whereas remaining five were designated FusC 2 thru FusC 6 (**Table S2**). These protein sequences ranged roughly in three different lengths: ∼200 amino acids for FusC 3, ∼350 for FusC 4 and 6, and ∼750 amino acids for FusC, FusC 2 and FusC 5. Upon examination, the loci around FusC genes 2 thru 6 were not arranged in as discernable tripartite-pump operons, like FusC, although some were adjacent to either a MFS transporter protein or genes encoding amino acid permeases. To test whether these genes are regulated by FA addition, we performed RT-qPCR on RNA isolated from Bp82 cultures induced with 1000 μM FA and without treatment. The transcription of *fusC* 2-6 genes showed little to no fold-change (0-2-fold; **Fig 6B**) when compared to non-treated samples, which contrasts with ∼40-fold difference in *fusC* transcription (**Fig. 4B**). Thus, we conclude that these *fusC* genes are not inducible by FA.

### Bucl8-pump does not contribute to the multidrug resistance (MDR) phenotype

Efflux pumps contribute to MDR in Gram-negative bacteria [11], including *Burkholderia* species [8], and are often polyspecific [51]. A study in *S. maltophilia* concluded that an FA efflux pump did not extrude the antimicrobials tested [50]. Here, we assessed changes in resistance/ susceptibility levels between Bp82 and Bp82Δ*bucl8-fusE*, and JM109 and JM109::525 or JM109:529 against variety of antimicrobials.

In the clinical laboratory setting, the *Burkholderia* failed to grow in commercial medium, and therefore only the *E. coli* data were generated. Overall, there was not a significant increase in resistance to any of the antibiotics tested; JM109::525/529 showed only increased resistance to the β-lactam antibiotics, which was associated with the resistance gene present on the inserted plasmid. A disc diffusion test, including ampicillin, ciprofloxacin, levofloxacin, tobramycin, gentamicin, tetracycline, doxycycline, and trimethoprim-sulfamethoxazole, resulted in similar zones of inhibition for both Bp82 and Bp82Δ*bucl8-fusE* cultures, as well as *E. coli* JM109 and JM109::525/529, again with the exception of the plasmid-derived β-lactam resistance determinate.

Microarray data comparing the effect of 84 growth conditions on *B. pseudomallei* transcriptome showed that chloramphenicol (CHL), which contains an aromatic ring in its structure, induced *bucl8* expression, thus, implying CHL might be a substrate for Bucl8-associated pump [52]. Here, we determined the CHL-MICs of our *B. pseudomallei* and *E. coli* strains using a growth assay on the LA medium; however, the MIC for all the strains was the same (8 µg/mL; **Fig 6A**). In addition, the exogenous CHL at 8 µg/mL or 4 µg/mL concentrations did not significantly induced the transcription of *bucl8*-associated genes (not shown). Thus, our results indicate the Bucl8-associated pump is not needed for CHL resistance in *B. pseudomallei* [49].

### Deletion of Bucl8-pump components affects cell growth

Efflux pumps extrude a variety of compounds that are toxic to the cells and play physiological functions linked to pathogenesis [12]. We observed the growth of the BpΔ*bucl8-fusE* mutant was considerably reduced than that of the parent Bp82 and did not reach the same OD_600_ in the stationary phase (**Fig 6C**). CFU for Bp82 increased by approximately four logs, while the mutant increased by two logs from hour 0 to 12. (**Fig 6D**). These results suggest that the pump components are needed for normal growth physiology under laboratory conditions in rich medium.

## Discussion

The protein Bucl8 was previously predicted to be the outer membrane in *B. pseudomallei* and *B. mallei*. Comparative genomics studies between *B. mallei* and *B. pseudomallei* have suggested that conserved genes between the species are likely critical for host-survival, while genes useful for saprophytic life-style and adaptability were selected against [6]. The presence of the *bucl8* genes, in particular the acquisition and conservation of the extracellular Bucl8-CL-Ct domain, in *B. pseudomallei* and *B. mallei* suggests that these genes are selected for because they are useful for bacterial survival in both the environment and in the host. Here, we carried out structure-function studies of the Bucl8 protein and associated locus in *B. pseudomallei* and *B. mallei* in order to elucidate the role of Bucl8 and its associated pump components in antimicrobial resistance, ligand binding, and cell physiology.

In the absence of hydroxyprolines that stabilize the triple helical structure of mammalian collagen, bacterial collagens adopt alternative stabilization mechanisms to form stable triple helices [53]. While some prokaryotic collagens utilize a variety of GXY-repeat types, such as streptococcal collagen-like proteins Scl1 and Scl2 [54], others possess a limited number of triplets, including *Bacillus* Bcl proteins [55, 56]. The CL regions of various Bucl proteins utilize relatively few distinct triplet types [13]. An extreme case is the Bucl8-CL region, which is exclusively made of a rare repeating (GAS)_n_ sequence. Our results are consistent with studies of triple helix propensity based on host-guest peptide studies, showing reasonable propensities of (GAS)_n_ triplets to form triple helical structures. The Tm value of (GAS)_n_ tripeptide unit in a triple helix is 33.0°C, compared to 47.3°C of (POG)_n_ tripeptide (O is hydroxyproline), although, the physical anchoring of a CL domain increases Tm by additional 2°C [57]. This relatively low Tm may suggest structural flexibility of the Bucl8 extracellular domain under physiological conditions, thus, allowing efflux pump for dual function.

Our laboratory and others have shown that bacterial collagen-like proteins participate in pathogenesis via a variety of functions, including immune evasion, cell adhesion, biofilm formation, and autoaggregation [43–45, 58]. Here, we report that the recombinant rBucl8-CL-Ct polypeptide binds to fibrinogen significantly better than rBucl8-Ct polypeptide. A similar phenomenon was recently reported for Scl1, where the effective binding to fibronectin, directly mediated by the globular V domain, required the presence of adjacent Scl1-CL domain [28]. Fibrinogen is a circulating glycoprotein that is involved in blood clotting and promoting wound healing [59]; we do not know the location of Bucl8 binding site on this multidomain protein. In the scope of pathogenesis, some Gram-negative and Gram-positive bacteria use fibrinogen for biofilm formation and bacterial adhesion. For example, fibrinogen-binding factors and clumping-factors of *Staphylococcus aureus* have been shown to increase adherence and virulence [60–62]. *B. pseudomallei* and *B. mallei* both cause cutaneous infections that lead to lesions and nodules, thus binding to wound factors could increase colonization. In addition, it is likely that unidentified ligand(s), other than fibrinogen, may exist in the environment to support a saprophytic lifestyle of *B. pseudomallei*.

*bucl8*-operon expression is regulated by a LysR-type transcriptional regulator, designated here as FusR_LysR_. LysR-type family regulators are the most abundant class of the prokaryotic transcriptional regulators that monitor the expression of genes involved in pathogenesis, metabolism, quorum sensing and motility, toxin production, and more physiological and virulence traits [49]. LysR proteins are tetrameric and consist of two dimers that bind and bend the DNA within promoter regions, thus, affecting the gene transcription. After the co-inducer binds to the LysR dimers, the DNA is relaxed, allowing one dimer to come into contact with the RNA polymerase to form an active transcription complex. In this study, the FusR binding sites were identified within the intergenic promoter region between *bucl8* and *fusR* in *B. pseudomallei* and *B. mallei*. Thus, we hypothesized that FA can act as a co-inducer for the *bucl8*-operon.

We show that exogenous fusaric acid (FA) induces the transcription of the *fusR-bucl8-fusCD-fusE* operon, therefore, confirming Bucl8 is a component of a previously unreported FA-inducible efflux pump in *B. pseudomallei* and *B. mallei*. Similarly, an inducible FA tripartite efflux pump, encoded by *fuaABC* operon, was identified in another soil saprophyte *S. maltophilia* [50]. However, the gene/protein arrangement, *e.g.* sequence orientation and length, places the *bucl8* operon within clade III of a phylogenetic tree of predicted FusC-associated operons, while *fuaABC* operon is in clade IV [37]. In addition to FA, the FA-derivative pHBA also induced the expression of the *bucl8* operon. Interestingly, although the genes and intergenic regions are highly similar, transcription of *fusR* and *bucl8* in FA-induced *B. mallei* culture is considerably reduced compared to *B. pseudomallei*. Likewise, the MIC levels for FA and pHBA were lower in *B. mallei*, although the *bucl8* loci are conserved between *B. pseudomallei* 1026b and *B. mallei* ATCC23344. There may be other factors affecting transcription, such as additional regulatory circuits for processing FA and similar compounds in both organisms. For similarity, another efflux pump in *B. pseudomallei*, BpeEF-OprC, is regulated by two highly similar LysR-type transcriptional regulators, BpeT and BpeS [63]. Further studies are needed to identify if there are other regulators or environmental stress/factors that could be affecting upstream/downstream targets.

Efflux systems are categorized into families by their structure – including their composition, conserved domains, and number of transmembrane spanning regions – as well as by their energy source and substrates. In *Fusarium*, the synthesized intracellular FA is extruded by a predicted MFS-type transporter FUBT [64], however based on the number of amino acids present and transmembrane helices, FusC is not likely a MFS transporter. Only ABC or RND systems regularly form tripartite complexes. It is not known whether the Bucl8 pump relies on ATP hydrolysis to transport FA, but the associated FusC_IMP_ transporter does not contain an ATP-binding domain, therefore, it is an unlikely an ABC-transporter. Phylogenetic analysis of bacterial efflux systems implied that FuaABC tripartite FA efflux pump in *S. maltophilia* forms a separate branch from other bacterial efflux pump families, branching off between the ABC and RND families [50]. Thus, we construe that the Bucl8 associated efflux pump is RND-like.

Here, we adopted the gene designation proposed by Crutcher *et al.*, which also includes a fourth pump component, a small polypeptide DUF, for the Bucl8-associated tetrapartite efflux system. This situation might be more common among known tripartite efflux pumps than currently acknowledged; for example, a small polypeptide YajC is an inner membrane component of a well-recognized “tripartite” RND system AcrAB-TolC [65]. Another known tetrapartite RND efflux system is the CusCFBA complex, which transports heavy metals copper and silver [66]. In this system, the small CusF component serves as a periplasmic metal-binding chaperone, which hands over the metal-ion substrate to the IMP transporter [67, 68]. The precise cellular location and function of FusD_DUF_ protein is not known at present.

Early studies reported FA-detoxification genes found in *Burkholderia* (formerly *Pseudomonas*) *cepacia* and *Klebsiella oxytoca* [15, 16], which were attributed to FA resistance. More recent work identified a tripartite FA efflux pump, FuaABC, in *Stenotrophomonas maltophilia* [50], while other work distinguished a large number of the phylogenetically related FusC-type proteins, conferring FA resistance, in numerous Gram-negative bacterial species [37]. Not all FusC proteins were predicted components of FA efflux pumps; however they were assumed to be contributing to high levels of FA resistance in some species, including *Burkholderia*. Crutcher *et al*. reported positive correlation between the number of putative FusC proteins in bacterial genomes and the level of resistance to FA; for example, *Burkholderia cepacia*, harboring six predicted FusC protiens, had a FA-MIC of ≥ 500 μg/mL, whereas *Burkholderia glumae* had two FusC proteins and a FA-MIC of 200 μg/mL [37]. Strains with 0-1 *fusC* genes were sensitive to FA with MIC <50 µg/mL. We also observed that our Bp82Δ*bucl8-fusE* mutant retained 100 μg/mL residual resistance to FA. Through transcriptional analysis, we found that the five *fusC*/FusC genes/proteins outside of the Bucl8-operon showed little to no induction, indicating that the Bucl8 pump is the main contributor to FA resistance in *B. pseudomallei*.

The multidrug resistance in *B. pseudomallei* is substantially attributed to previously studied RND efflux pumps BpeAB-OprB, AmrAB-OprA, and BpeEF-OprC. At the same time, little is known about the role of FA pumps in resistance against clinically used drugs. In our studies, we assessed the Bucl8-pump’s role in multidrug resistance in two ways: (i) we compared the spectrum of resistance between parental strain Bp82 and Bucl8-pump deletion mutant, and (ii) we expressed the *bucl8*-operon in a sensitive *E. coli* strain. Although MICs for FA changed as predicted, deletion of the Bucl8-pump did not affect MIC values for the clinically-used drugs. This result is comparative to an FA pump in *S. maltophilia*, which did not determine the resistance to a large panel of therapeutics tested [50]. At the same time, a different study in the same organism showed that deletion of the *pcm-tolCsm* operon, encoding a different efflux pump, resulted in decreased MICs for several antimicrobials of diverse classes (β-lactams, chloramphenicol, quinolone, tetracycline, aminoglycosides, macrolides), and also decreased FA resistance [69]. Microarray data suggested *bucl8* expression was upregulated in the presence of chloramphenicol [52] and deletion of the *tolCsm* in *S. maltophilia* resulted in decreased resistance to both CHL and FA [69]. Both CHL and FA harbor aromatic rings in their structures, however, our investigations did not detect *bucl8*-operon induction by CHL nor changes in CHL resistance levels in Bp82Δ*bucl8-fusE* mutant or complemented *E. coli*.

The decrease in bacterial growth of the Bp82Δ*bucl8-fusE* mutant suggests that the Bucl8-pump may be involved in modulating essential cellular stresses, both in the environment and in infected human host [12]. Limited studies show that FA repressed quorum sensing genes, expression of stress factors, secretion of siderophores, production of anti-fungal metabolites, and iron uptake [70–73]. Additionally, Bucl8 pump may be involved in a transport of aromatic carboxylic acid compounds and act as a pHBA-metabolic efflux valve [38]. Further investigation will be needed to determine what cellular processes are associated with the Bucl8-pump.

In summary, we conclude that Bucl8 is a component of a previously unreported tetrapartite efflux system that is involved in FA resistance and cell physiology. We have demonstrated that the extracellular Bucl8-CL domain forms the prototypic collagen triple-helix, while the extracellular Bucl8-CL-Ct portion is capable of binding to fibrinogen. Further studies will investigate what role fibrinogen binding plays in pathogenesis. While the Bucl8-pump is likely not be involved in the MDR phenotype of *Burkholderia*, we have identified FA and pHBA as inducible substrates of the pump and will continue to investigate metabolite analogs that may affect cell function. Importantly, the growth of the Bucl8-pump deletion mutant was significantly affected even in the absence of FA and pHBA. By characterizing the Bucl8-associated efflux system, we can advance therapies and strategies for combating these pathogens, including developing pump inhibitors, targeting transport mechanisms, or identifying potential surface-exposed vaccine targets derived from Bucl8.

## Acknowledgements

We would like to thank Heath Damron for providing us pUC18T-mini-Tn7T-Tp vector, Alfredo Torres for providing *B. mallei* CLH001, Shelby Bradford for her assistance with initial experiments, and Paul Feustel for advice with statistical analyses.

Opinions, interpretations, conclusions, and recommendations are those of the authors and are not necessarily endorsed by the US Army.

