## Supplementary material for "*Burkholderia* collagen-like protein 8, Bucl8, is a unique outer membrane component of a tetrapartite efflux pump in *Burkholderia pseudomallei* and *Burkholderia mallei*": Table S

Table S1. Primers

| **Target** | **Primer name** | **Primer sequence** |
| --- | --- | --- |
| *fusR* | pSL522-ApaI-F | 5’-GAAGGGCCCATGCTTGCGCATCCGG-3’ |
|  | pSL522-HindIII-R | 5’-GGTAAGCTTCGGGCATCACGCGCACG-3’ |
|  | BurkhLysR-2F | 5’-GTTCGTCCGCGTGGTCGATG-3’ |
|  | BurkhLysR-2R | 5’-CGCAAGTGCGCCTCGAGATC-3’ |
| *bucl8* | Bucl8-1F | 5’-CTCGTATGAAGAGGCGATCC-3’ |
|  | Bucl8-3F | 5’-CTACGCGCTCCTCGACATCGCGC-3’ |
|  | Bucl8-3R | 5’-TGCGTGCCGATGCCCGCGCGCA-3’ |
| *fusCD* | BurkhFusBCD-1F | 5’-GTGGCTCTATCTCGCGAAGGCGC-3’ |
|  | BurkhFusBCD-1R | 5’-GCGGCTGCATCACGATGAACACGG-3’ |
| *fusE* | BurkhFusE-1F | 5’-CAGCCGTCATCCTGATCGTCGCG-3’ |
|  | BurkhFusE-1R | 5’-CGGCGCGACGTTGACGATCTCC-3’ |
| *tar* | pSL523-ApaI-2F | 5’-CGACTTGCGCTTGCCGCCGGGCCCTTG-3’ |
|  | pSL523-ApaI-2R | 5’-GAAGGGCCCGCGACGAGCATGGGGCAAC-3’ |
|  | BurkhTar-1F | 5’-CGCACGATGGACGAGGTCGTGC-3’ |
|  | BurkhTar-1R | 5’-CCCGCGCTCTGCTCACTCGACG-3’ |
| Plasmids | pMo130-MCSI-F | 5’-GCTCACATGTTCTTTCCTGCG-3’ |
|  | pMo130-MCSI-R | 5’-CCCGGTCGCATTACACCTTTG-3’ |
|  | pSL520-F | 5’-CACGGATCCTCGACTGC-3’ |
|  | pSL520-R | 5’-CAAGCTTTAGCGAGCTGCA-3’ |
|  | pSL521_1F | 5’-GAGGAGAAATTAACTATGAGAGGATCG-3’ |
|  | pSL521_2R | 5’-AGCTAATTAAGCTTTAGCGAGCTG-3’ |
|  | pQE30-F | 5’-CACCTGACGTCTAAGAAACCATTAT-3’ |
|  | pQE30-2R | 5’-TCTCCATTTTAGCTTCCTTAGCTCC-3’ |
| FUSC | FusC 2-F | 5’-TGTCGCTCATCGTCGTCTA-3’ |
|  | FusC 2-R | 5’-AGCGGCGTGAATTTCTCTT-3’ |
|  | FusC 3-F | 5’-GATCGTGACGGCGATCATCTG-3’ |
|  | FusC 3-R | 5’-GGAAACAGCAGCACGATTGTC-3’ |
|  | FusC 4-2F | 5’-CCACGGCGATAACGAGATCGC-3’ |
|  | FusC 4-2R | 5’-CACGAGTAGGTCGCATACAGC-3’ |
|  | FusC 5-2F | 5’-CGATGAGCGGGATGGTGCTC-3’ |
|  | FusC 5-2R | 5’-CGCGACCCACAGCGTGATGC-3’ |
|  | FusC 6-F | 5’-CGTCGATCGCGACGGTATCG-3’ |
|  | FusC 6-R | 5’-GCACGACAGCGACAGAAAGCC-3’ |
| 16s | 16s rRNA-F | 5’- GGCTAATACCCGGAGTGGA-3’ |
|  | 16s rRNA-R | 5’- CTAGCCTGCCAGTCACCAA-3’ |

Table S2. Genes and associated identification numbers of FusC loci

|  | **Bp 1026b** | | **Bp K96243** | | **Bm ATTC 23344** | |
| --- | --- | --- | --- | --- | --- | --- |
| **Gene** | **Locus tag** | **Protein ID** | **Locus tag** | **Protein ID** | **Locus tag** | **Protein ID** |
| *fusC 2* | BP1026B_RS11380 | [WP_004552879.1](https://www.ncbi.nlm.nih.gov/protein/490656881) | BPS_RS06755 | [WP_004534567.1](https://www.ncbi.nlm.nih.gov/protein/490669577) | DM55_RS13405 | [WP_004193403.1](https://www.ncbi.nlm.nih.gov/protein/490297964) |
| *fusC 3* | BP1026B_RS12100 | [WP_004198110.1](https://www.ncbi.nlm.nih.gov/protein/490302733) | BPS_RS05900 | [WP_004198110.1](https://www.ncbi.nlm.nih.gov/protein/490302733) | DM55_RS12800 | [WP_004198110.1](https://www.ncbi.nlm.nih.gov/protein/490302733) |
| *fusC 4* | BP1026B_RS21370 | [WP_004187681.1](https://www.ncbi.nlm.nih.gov/protein/490292089) | BPS_RS21435 | [WP_004187681.1](https://www.ncbi.nlm.nih.gov/protein/490292089) | DM55_RS20330 | [WP_004202241.1](https://www.ncbi.nlm.nih.gov/protein/490307172) |
| *fusC 5* | BP1026B_RS22725 | [WP_004539022.1](https://www.ncbi.nlm.nih.gov/protein/490674193) | BPS_RS22800 | [WP_004539022.1](https://www.ncbi.nlm.nih.gov/protein/490674193) | DM55_RS24535 | [WP_004195004.1](https://www.ncbi.nlm.nih.gov/protein/490299590) |
| *fusC 6* | BP1026B_RS28905 | [WP_004552079.1](https://www.ncbi.nlm.nih.gov/protein/490687587) | BPS_RS29185 | [WP_004525018.1](https://www.ncbi.nlm.nih.gov/protein/490660028) | DM55_RS17915 | [WP_004190560.1](https://www.ncbi.nlm.nih.gov/protein/490295011) |

Data were retrieved from NCBI for *B. pseudomallei* 1026b, *B. pseudomallei* K96243, and *B. mallei* ATTC 23344 reference genomes. Proteins were labeled as FUSC family protein.

**Table S3. gBlock inserts for construction of recombinant proteins**

| Construct/ protein | Amino Acid Sequence | Nucleotide Sequence |
| --- | --- | --- |
| pSL520/ rBucl8-Ct | MRGSHHHHHHGSSTAGASATASASAAGHAPAGAAAPASPAGIRAAASARASMPAPAAAATAPAFASPVAGASTPMPAATAAARAAR | CACGGATCCTCGACTGCGGGAGCATCTGCAACTGCAAGCGCAAGCGCGGCTGGCCATGCACCGGCTGGCGCGGCAGCTCCAGCGTCACCAGCTGGCATTCGCGCTGCTGCTTCGGCGAGAGCGTCTATGCCGGCACCAGCCGCAGCGGCAACTGCGCCTGCCTTTGCATCACCGGTGGCTGGTGCTTCGACTCCGATGCCTGCAGCAACCGCAGCTGCTCGTGCAGCTCGCTAAAGCTTG |
| pSL521/ rBucl8-CL-Ct | MRGSHHHHHHGSGASGASGASGASGASGASGASGASGASGASGASGASGASGASGASGASGASGASSTAGASATASASAAGHAPAGAAAPASPAGIRAAASARASMPAPAAAATAPAFASPVAGASTPMPAATAAARAAR | GAGGAGAAATTAACTATGAGAGGATCGCATCACCATCACCATCACGGATCCGGTGCTTCGGGTGCTTCGGGTGCCTCGGGTGCCTCGGGTGCCTCGGGTGCCTCGGGTGCCTCGGGTGCCTCGGGTGCCTCGGGTGCCTCGGGTGCCTCGGGTGCCTCGGGTGCCTCGGGTGCCTCGGGTGCCTCGGGTGCCTCGGGTGCCTCGGGTGCCTCGGGTGCCTCGGGTGCCTCGGGTGCTTCGTCGACTGCGGGAGCATCTGCAACTGCAAGCGCAAGCGCGGCTGGCCATGCACCGGCTGGCGCGGCAGCTCCAGCGTCACCAGCTGGCATTCGCGCTGCTGCTTCGGCGAGAGCGTCTATGCCGGCACCAGCCGCAGCGGCAACTGCGCCTGCCTTTGCATCACCGGTGGCTGGTGCTTCGACTCCGATGCCTGCAGCAACCGCAGCTGCTCGTGCAGCTCGCTAAAAGCTTAATTAG |
